# Single-channel in-Ear-EEG detects the focus of auditory attention to concurrent tone streams and mixed speech

**DOI:** 10.1101/094490

**Authors:** Lorenz Fiedler, Malte Wöstmann, Carina Graversen, Alex Brandmeyer, Thomas Lunner, Jonas Obleser

**Affiliations:** Department of Psychology, University of Lübeck, Germany; Max Planck Research Group “Auditory Cognition,” Max Planck Institute for Human Cognitive and Brain Sciences, Germany; Eriksholm Research Centre A/S, Oticon, Denmark

**Keywords:** BCI, EEG, Hearing aid, auditory, selective attention, forward model

## Abstract

**Objective:** Conventional, multi-channel scalp electroencephalography (EEG) allows the identification of the attended speaker in concurrent-listening (“cocktail party”) scenarios. This implies that EEG might provide valuable information to complement hearing aids with some form of EEG and to install a level of neuro-feedback.

**Approach:** To investigate whether a listener’s attentional focus can be detected from single-channel hearing-aid-compatible EEG configurations, we recorded EEG from three electrodes inside the ear canal (“in-Ear-EEG”) and additionally from 64 electrodes on the scalp. In two different, concurrent listening tasks, participants (n = 7) were fitted with individualized in-Ear-EEG pieces and were either asked to attend to one of two dichotically-presented, concurrent tone streams or to one of two diotically-presented, concurrent audiobooks. A forward encoding model was trained to predict the EEG response at single EEG channels.

**Main results:** Each individual participants’ attentional focus could be detected from single-channel EEG response recorded from short-distance configurations consisting only of a single in-Ear-EEG electrode and an adjacent scalp-EEG electrode. The differences in neural responses to attended and ignored stimuli were consistent in morphology (i.e., polarity and latency of components) across subjects.

**Significance:** In sum, our findings show that the EEG response from a single-channel, hearing-aid-compatible configuration provides valuable information to identify a listener’s focus of attention.

## 1. Introduction

In multi-talker situations, hearing-aid users find it difficult to comprehend the attended conversational partner against background noise (i.e. cocktail party problem, Cherry 1953). Part of this problem might be due to the fact that the hearing aid is lacking the explicit information which sound source the listener wants to listen to. The investigation of neural speech-tracking (for a methods review, see Wöstmann *et al* (2016)) using Electroencephalography (EEG) and identification of the attended speaker in multitalker scenarios from multi-channel scalp-EEG (Mirkovic *et al* 2015, O’Sullivan *et al* 2015) has demonstrated that EEG could feasibly inform future hearing aid algorithms about a listener’s focus of attention. Information about the focus of attention would allow hearing aids for example to adapt noise suppression algorithms or to align directional microphones to the attended sound source (Mirkovic *et al* 2016, Van Eyndhoven *et al* 2016).

The implementation of EEG into comparably small hearing aids allows the attachment of only few electrodes at restricted positions inside the ear canal (Mikkelsen *et al* 2015, Bleichner *et al* 2015) or around the ear (Mirkovic *et al* 2016, Debener *et al* 2015). Since EEG responses quantify the potential difference between a signal electrode and a reference potential, at least two electrodes are required to measure the EEG. The position and distance as well as the orientation of the two electrodes mainly determines, in how far relevant and irrelevant electrophysiological and external sources will be captured, respectively. Due to the limited number of channels in such a hearing-aid-compatible configuration, established offline methods of EEG-signal enhancement such as independent component analysis relying on covariance of multiple, whole scalp covering electrodes (Makeig *et al* 2004) are not applicable.

An established method to extract auditory evoked potentials (AEP) is based on multiple time-locked presentations of identical stimuli and the subsequent averaging of the measured EEG time-domain signal (Rockstroh *et al* 1982). Using this method, it has been shown that the AEP can be extracted from the potential difference between in-Ear-EEG electrodes and adjacent scalp-EEG electrodes (Fiedler *et al* 2016, Mikkelsen *et al* 2015, Bleichner *et al* 2015). For the presentation of continuous, non-repeating speech, averaging across multiple trials is not applicable (for review see Wöstmann *et al* 2016). Thus, a method to estimate a response evoked by continuous speech is needed. Importantly, the quasi-rhythmic fluctuations of the speech signal’s broad-band temporal envelope have recently been reconstructed successfully from Magnetoencephalography (MEG) (Ding and Simon 2012) and EEG (O’Sullivan *et al* 2015, Mirkovic *et al* 2015) using linear models. Despite some remaining ambiguities as to the signal features that do get actually encoded in the neuro-cortical signal (see e.g. Ding and Simon 2014), a main finding here is that the attended-speaker signal attains a dominant representation in the measured neural signal.

In sum, recent scalp-EEG research has established the feasibility to infer on a listener’s attentional focus from EEG very generally. In this present study, however, the overriding goal is to examine single-channel in-Ear-EEG configurations that possibly could be part of a hearing aid. To this end, we focus our analyses on single-channel electrode configurations consisting of an in-Ear-EEG and a scalp-EEG electrode close to the ear only, to allow future smooth integration with extant hearing-aid systems (Lunner and Gustafsson 2014). We employ estimation of a forward (i.e., encoding) model since we focused on the encoding of onsets in the broad-band temporal envelope and the prediction of the to-be-expected EEG-signal at single EEG channels. Furthermore, we avoided any methods of artefact rejection such as independent component analysis or trial rejection. This approach allows us to presume that the same results could have been achieved by solitarily recording the respective channel by attaching only two electrodes.

The resulting data from two challenging, cocktail-party-like listening paradigms demonstrate that, on the single-participant level, we are able to accurately infer a listener’s attentional focus from a single-channel EEG setup consisting of electrodes in and around the ear.

## 2. Methods

### 2.1 Participants

Eight subjects were enrolled in the study (aged 23, 25, 28, 29, 39, 41, 43 and 49; 4 males). Each participant was provided with individually fitted ear molds. Each ear mold was equipped with three in-Ear-EEG electrodes (Fiedler *et al* 2016).

Five of the subjects were native Danish speakers, while two were French and one was a German native speaker. All reported normal hearing and no histories of neurological disorders. Participants gave informed consent. Procedures were in accordance with the Declaration of Helsinki and approved by the local ethics committee of the University of Leipzig Medical faculty. All subjects participated in the oddball task, while only the five native Danish speakers participated in the audiobooks task (aged 29, 39, 41, 43 and 49, 3 males). For both tasks, the recording from one of the Danish subjects had to be discarded due to invalid in-Ear-EEG data, as the device did not remain in place during recordings.

Note that the comparably low number of subjects is due to the fact that the in-Ear-EEG devices are in a prototype stadium and can’t be manufactured in high quantities. However, all results presented are based on rigorous levels of statistical significance in the single subject.

### 2.2 Stimuli and Tasks

We implemented two experimental paradigms in order to investigate whether neural responses for two concurrent auditory streams can be extracted from in-Ear-EEG and whether such responses can predict which out of two streams is being attended.

First, we implemented a non-speech, two-stream, dichotic tone paradigm, in close analogy to Lakatos *et al* (2012), hereafter called oddball task. Two dichotically presented (i.e., left vs right ear) concurrent streams of 100-ms tones (with a sawtooth carrier waveform) were presented for one minute. On each trial, the two streams differed in tone repetition rate (1.4 vs. 1.8 Hz) and pitch (410 vs. 610 Hz). 10-15 % of the tones occurred as oddballs (1/4 tone pitch deviation) in both streams. Participants were asked to either attend to the stream presented on the left or right ear and to press a button with their right hand as soon as they heard an oddball in the attended stream. In total, 40 trials of one minute length were presented (Figure 1A). All stimulus manipulations, repetition rate (1.4 vs 1.8 Hz), pitch (410 vs 610 Hz), and attention (left vs right) were counterbalanced across trials.

**Figure 1:**
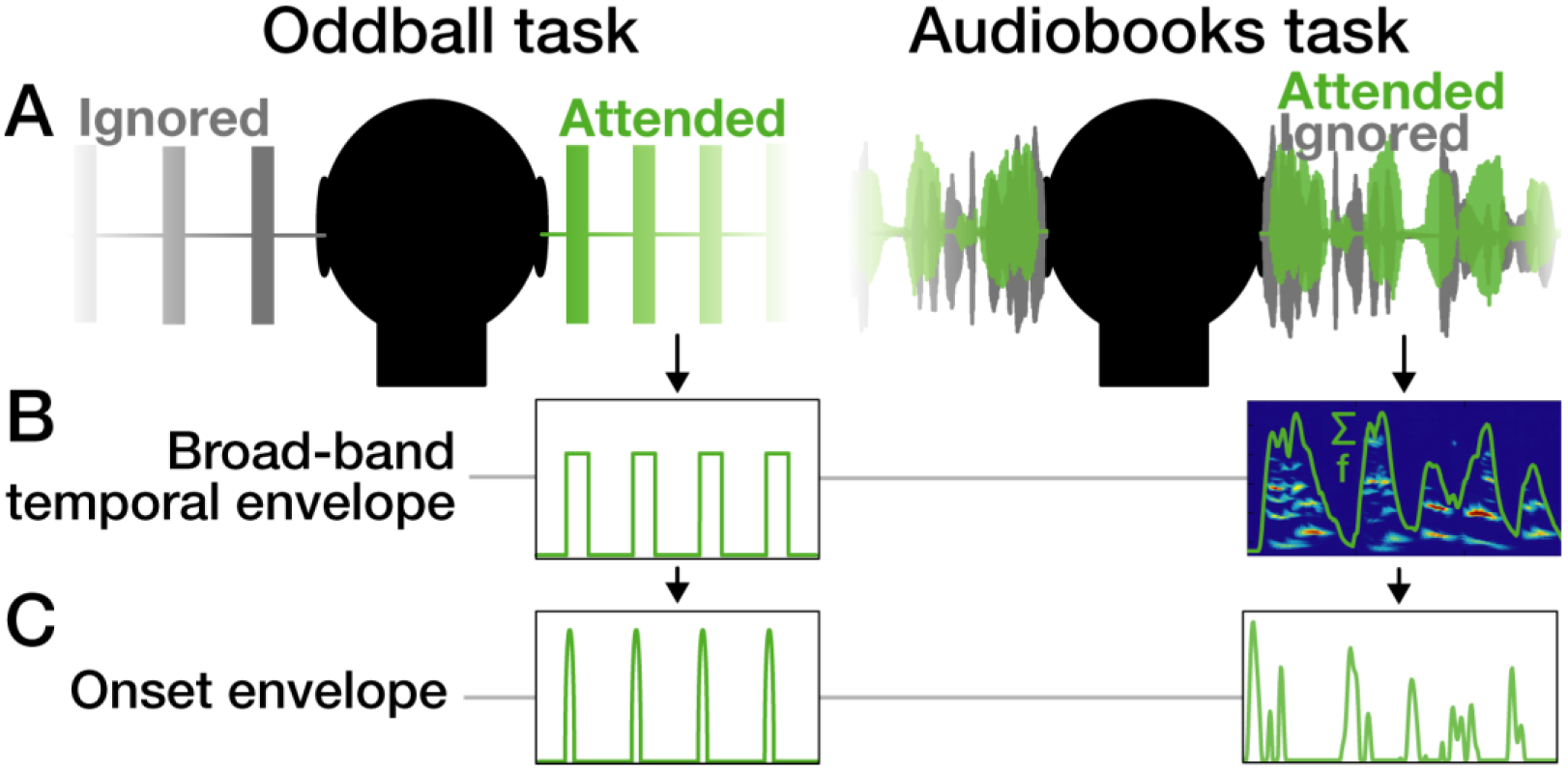
Design and Onset envelope extraction. **A)** Exemplary stimulus waveforms show the spatial separation of target (green) and distractor (grey) stimuli in both tasks. In the oddball task, two streams of 100-ms tones differing in repetition rate and pitch were presented. Subjects were asked to attend to the left or the right stream and press a button as soon as they heard an oddball (pitch deviation) in the attended stream. In the audiobooks task, two Danish audiobooks spoken by a female and male speaker were presented. The identical mixture of both speakers was presented on both ears (diotic). Subjects were asked to attend either the female or the male voice. **B)** In the oddball task, the broad-band temporal envelope was captured from the stimulus-waveforms directly. In order to capture the broad-band temporal envelope from the audiobooks, an auditory time-frequency representation was summed up across its spectral sub-bands. **C)** The onset envelope was obtained by computing the first derivative of the broad-band temporal envelope and subsequently zeroing values smaller than zero (half-wave rectification).

The second paradigm was a two-stream, continuous-speech paradigm, hereafter called audiobooks task. Emulating typical challenging listening scenarios, we presented a mixture of two concurrent audiobooks to both ears (i.e., diotic presentation without any spatial cues; figure 1A). The stimuli were two different Danish works of fiction spoken by a female (F. Marryatt, *Children of the forest)* and a male speaker (E. A. Poe, *A Descent into the Maelström)*, with matched long-term root-mean-squared (rms) sound intensity. Each exemplar of one-minute mixtures was presented twice in succession. Counterbalanced across trials, subjects were asked to either attend to the male voice first and second to the female voice or vice versa. In total, 60 trials of such one-minute mixtures were presented.

### 2.3 EEG-Data Acquisition and Preprocessing

Sixty-four–channel scalp-EEG was recorded alongside in-Ear-EEG using a *BioSemi ActiveTwo* amplifier (*Biosemi*, Netherlands). In-Ear-EEG electrodes were connected to the auxiliary inputs of the *ActiveTwo* amplifier via pre-amplifiers identical to the ones used for scalp-EEG electrodes. EEG data were recorded with a sampling rate f_s_ = 2048 Hz. Please find more details about the recording procedure in Fiedler *et al* (2016).

Data were preprocessed using both the *fieldtrip toolbox* (Oostenveld *et al* 2011) for *Matlab* (*MathWorks, Inc.)* and custom-written code. The continuous EEG data recorded during the oddball task were highpass-filtered at f_c_ = 1 Hz and lowpass-filtered at f_c_ = 15 Hz. The continuous EEG data recorded during the audiobooks task were highpass-filtered at f_c_ = 2 Hz and lowpass-filtered at f_c_ = 8 Hz according to O’Sullivan *et al* (2015). In order to compensate phase shifts, data were filtered both forward and backward using Hamming-window FIR filters with orders N = 3f_s_/f_c_. Subsequently, all data were downsampled to 125 Hz to match the sampling rate of the onset envelopes (see below).

After an initial inspection of the event-related potential (ERP) between in-Ear-EEG electrodes and Cz, we encountered the issue of not all in-Ear-EEG electrodes keeping proper conductance across the whole experiment. Thus, for each ear canal, only the electrode showing minimal standard deviation across trials in the ERP summed up between 0 and 500 ms relative to tone-onsets was selected for further analysis.

In order to evaluate the potential difference between in-Ear-EEG electrodes and scalp-EEG electrodes, we created two datasets for each participant, one with all scalp-channels referenced to the priorly selected left in-Ear-EEG electrode and the other with all scalp-EEG channels referenced to the selected right in-Ear-EEG electrode.

### 2.4 Extraction of Onset Envelopes

Several approaches to extraction of the broad-band temporal envelope from a speech signal have been proposed (Biesmans *et al* 2016, Thwaites *et al* 2016). In case of the oddball task, the envelope was extracted by a direct calculation of the absolute values of the Hilbert-transform. In case of broad-band speech signals, the Hilbert transform is only a rough approximation and it has been shown that an intermediate step of extraction and subsequent summation of frequency sub-band envelopes increases the accuracy of detecting the attended speaker (Biesmans *et al* 2016). Thus, for the audiobooks task, we extracted the sub-band envelopes using NSL Toolbox (Ru 2001) for Matlab (Mathworks, Inc.), which resulted in a representation containing the envelopes of 128 frequency bands of uniform width on the logarithmic scale with center frequencies logarithmically spaced between 0.1 and 4 kHz (24 bands per octave). In order to obtain the broad-band temporal envelope, sub-band envelopes were summed up across frequency (figure 1B).

Furthermore, it has been proposed to transform the broad-band temporal envelope in order to extract salient increases of signal power (Hertrich *et al* 2012, Hambrook and Tata 2014). This method is based on the assumption that earliest time points of sensation that could evoke responses are tone or syllable onsets, respectively. It can be calculated by zeroing negative values (halfwave rectification) of the first derivative of the broad-band temporal envelope and results in a pulse-train-like series of peaks. Most salient peaks occur both at tone or syllable onsets (Fig 1C). This time-series will be called *onset envelope.* Recently, we have shown that the cross-correlation of the onset envelope and the EEG-signal results in estimations of the neural response similar to conventional ERPs obtained by multi-trial averaging (Fiedler *et al* 2016).

### 2.5 Training EEG response models

A schematic illustration of the approach to identification of the attended speaker is provided in Figure 2. In order to evaluate the performance in identification of the attended speaker at every single EEG channel, we first trained a model for each individual participant. The model is a linear mapping of the onset envelope onto the measured EEG signal.

**Figure 2:**
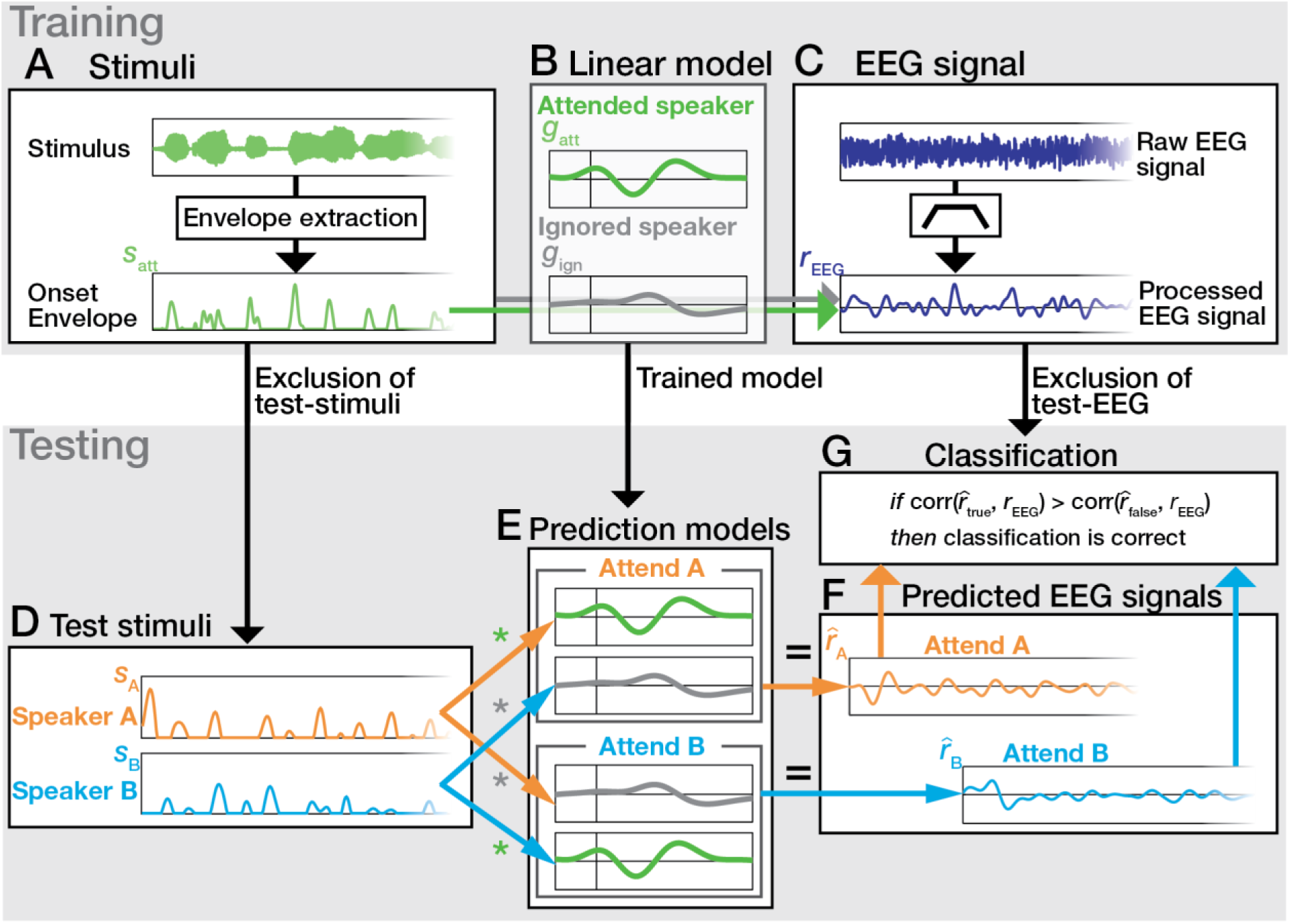
Identification of the attended speaker from single-channel EEG exemplary for audiobooks task. *Training:* After extraction of the onset-envelope (**A**) and preprocessing of the EEG signal (**C**), a linear forward model (**B**) is estimated for each trial and each speaker by concatenated stimulus and EEG signal of all other trials. *Testing:* The convolution of the onset envelopes of speaker A and B (**D**) with the trained prediction models (**E**) predicts to be expected EEG signals f_A_ and f_B_ with the labels *‘Attend A’* and *‘Attend B’,* respectively (**F**). **G**) If the predicted EEG signal labeled *true* (i.e., corresponds to the trial instruction) yields higher Pearson-correlation coefficient with the measured EEG-signal than the predicted EEG signal labeled *false* (i.e., is contrary to trial instruction), the classification is correct.

We used a well-established form of regularized regression (i.e., ridge regression; Hoerl and Kennard 1970) to train our model, as ridge regression has been shown to be applicable for predicting neurophysiological signals on the base of stimulus features (forward encoding model) (Santoro *et al* 2014, Lalor *et al* 2009) as well as reconstructing stimulus features from EEG signals (backward decoding model; O’Sullivan *et al* 2015, Mirkovic *et al* 2015). A *Matlab*-toolbox (*mTRF Toolbox*) is provided by Lalor (https://sourceforge.net/projects/aespa). As established above, the EEG signal should be independently predicted for every single EEG channel, which is, due to the implementation, inherent of forward modelling (Crosse *et al* 2016).

In detail, a single-channel encoding model *g* is the linear mapping of the onset envelope s onto the EEG signal *r*, which can be expressed as a convolution operation

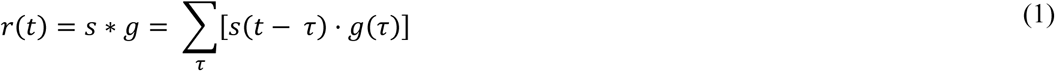

where *t* for *t* = 1,2,…,L is the sample index of both of the onset envelopes and the EEG signal with length L and τ for τ_min_, τ_min_+1,…, τ_max_ is the investigated sample-wise time lag between *s* and *r*. We investigated time lags (between the envelope and the EEG signal) ranging from −100 to 550 ms. In our design, we expect a difference in morphology of the response functions *g*_att_ *and g*_ign_ (figure 2B), which are models of the responses to the attended and the ignored stimulus onset envelopes *s*_att_ and *s*_ign_ (figure 2A). Moreover, we assume that the responses *r*_att_ and *r*_ign_ sum up and some noise *n* interferes (Zion Golumbic et al., 2013). Accordingly, we can express the measured EEG signal *r*_EEG_ (figure 2C):

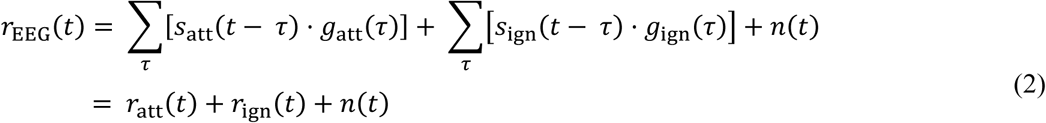

Since our goal was to estimate a response model including *g*_att_ *and g*_ign_ that minimizes the mean-squared error of the subsequent predicted EEG response *r*̂_EEG_, it can be obtained by the standard matrix operation in regularized regression,

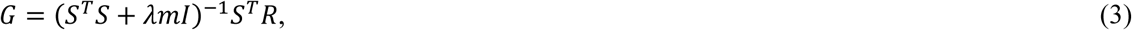

where S is an L-by-2T-matrix with its columns containing onset envelopes of both the attended *s*_att_ and ignored *s*_ign_ stimulus onset envelopes and their time-lagged replications. R is a column vector of length L containing the measured single channel EEG signal *r*_EEG_. The relative regularization parameter λ is first multiplied with m, the mean of the diagonal elements of S^T^S (Biesmans *et al* 2016). Second, it is multiplied with the identity matrix I and added to the covariance-matrix S^T^S. This regularization term λmI prevents overfitting (Crosse *et al* 2016), which appeared as high frequent artifacts in the to be estimated response models. The resulting matrix G contains the time-lag-wise response weightings *g*_att_ and *g*_ign_ for both the attended and ignored stimulus onset envelopes.

After an initial inspection of the response models, we decided to choose λ=10^2^. Please note that the greater λ is chosen, the more the term (S^T^S + λmI) converges to a multiple of the identity matrix, and the influence of covariance vanishes. This would lead to the same results as cross-correlation, which was also shown to be feasible for extracting neural responses (Kong *et al* 2014, Fiedler *et al* 2016), but doesn’t account for potential confounds caused by auto-correlation in the unregularized signal. Here we couldn’t observe a consistent benefit of regularization, because classification accuracy didn’t decrease by further increasing λ. However, in order to be consistent with the literature, we applied regression as stated above.

In line with former studies (O’Sullivan *et al* 2015, Mirkovic *et al* 2015), we decided to apply leave-one-out cross-validation. According to Biesmans *et al* (2016) we trained the prediction models by concatenating both the stimuli and EEG signal of all but the to-be-tested trial, before feeding it into (3). Thus, we obtained a prediction model for every single trial.

### 2.6 Testing EEG response models: Identification of the attended stream

In order to classify which of the streams a listener attended to, the former trial-wise trained models *g*_att_ and *g*_ign_ (figure 2B) were assembled to become two contrary prediction models (figure 2E). According to (1), the sum of the convolution of the onset envelopes *s*_A_ *and s*_B_ (figure 2D) and each response model (figure 2E) predicts an EEG signal, respectively. For both scenarios with the labels *Attend A* and *Attend B,* EEG signals *r*̂_A_ and *r*̂_B_ (figure 2F) were predicted:

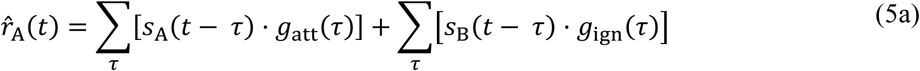

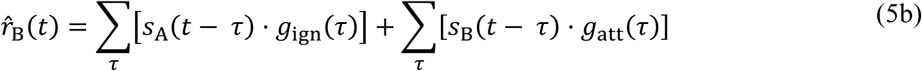

This operation can be expressed by matrix multiplication of the onset envelope matrix S and the response model matrix G:

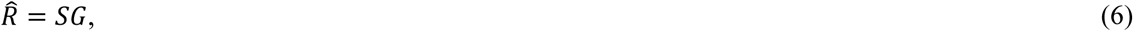

where *R̂* is a column vector containing the predicted EEG signal *r̂*_*A*_ or *r̂*_B_, respectively.

In order to estimate which of the predicted EEG signals (*r̂*_A_ vs *r̂*_B_) is most likely representing the trial instruction (attend A vs attend B), we calculated the Pearson-correlation coefficient of the predicted EEG signals (*r*̂_A_ and *r*̂_B_) and the measured EEG signal *r*_EEG_ respectively (L = 7500 samples, figure 2G). The predicted EEG signal that matched the to-be-attended stream (A vs B) was labeled *true,* the other one was labeled *false.* The classification was considered correct if the predicted EEG signal labeled *true* yielded greater (i.e., more positive) correlation than the EEG signal labeled false.

### 2.7 Goodness of fit

As a measure for the *goodness of fit,* we will refer to the correlation coefficient obtained from Pearson-correlation of the *true* prediction and the measured EEG signal. The greater this coefficient, the more of the measured EEG signal’s variability would be explained by the response model. Due to the fact that a convolution is a weighted sum and here the weights are the response models with positive or negative weights at certain time lags, the predicted EEG signals should have the same polarity as the measured EEG signal. Hence, the inspection of the correlation-coefficient’s magnitude (or square) wouldn’t be appropriate. Thus, a greater (i.e. more positive) correlation-coefficient indicates the *true* prediction.

### 2.8 Classification accuracy

By *classification accuracy* we will refer to the percentage of trials in which the predicted EEG signal labeled *true* yields higher correlation with the measured EEG-signal than the predicted EEG signal labeled *false.* For statistical analyses, both the correlation coefficients resulting from Pearson-correlation of the *true* and the *false* prediction with the measured EEG signal, respectively, were fisher-z-transformed and called z_true_ and z_false_. Considering the number of trials and the binary nature of the decision between two alternatives *Attend A* or *Attend B,* a single-subject chance level was defined at a level of significance α = 0.05 based on a binominal distribution (O’Sullivan *et al* 2015, Mirkovic *et al* 2016). This resulted in thresholds of 65% for the oddball task (40 trials) and 61.67% for the audiobooks task (60 trials).

## 3 Results

The main goal of this study was to identify the attended stimulus stream based on responses at single-channel EEG configurations consisting of one in-Ear-and one scalp-EEG electrode. To this end, we trained forward encoding models in order to predict EEG signals containing the predicted responses to both the attended and the ignored stimulus stream. Two alternative EEG signals representing the scenarios *Attend A* and *Attend B* were predicted. The prediction corresponding to the to-be-attended stream was called *true* and the other one *false. Goodness of fit* was quantified by Pearson-correlation coefficient of the *true* predicted and the measured EEG signal. For further statistical analyses, this coefficient was Fisher-z-transformed and called z_true_, whereas its counterpart z_false_ was equivalently computed by correlation of the false prediction and the measured EEG signal. Our approach to classification relies on the assumption that the true prediction better fits the measured EEG signal and thus leads to more positive correlation coefficients than the false prediction. Based on that, the percentage of correctly classified trials will be referred to as *classification accuracy.* All plots but the topographic maps are showing data from the exemplary configuration of FT7 referenced to the left in-Ear-EEG channel.

### 3.1 Response functions reveal consistent attention-related differences

Applying ridge regression to obtain a forward models is known to return response functions comparable to ERPs (Lalor *et al* 2009, Fiedler *et al* 2016). Beyond that, ridge regression can be applied on data measured during the presentation of continuous stimuli such as speech. According to (5), the aforementioned difference between the correlation coefficients z_true_ and z_false_ (see *below*) has to arise from differences between the response functions of the attended and ignored stimuli.

An inspection of the grand average response functions averaged across subjects in the dichotic oddball task (figure 3A) indicated that we extracted components equivalent to a P50-N100-P200 complex. The response functions (figure 3A) suggest an enhanced N100-equivalent component in responses to attended tones, which can be confirmed by the consistent differences of the responses to attended and ignored tones (Figure 3C). All subjects show a negative deflection in responses to attended tones at around 160 ms, while all but one of the subjects show a positive deflection in responses to attended tones at around 380 ms. The topographies of the differences at time lag of maximal deflections show a bilateral pattern.

**Figure 3:**
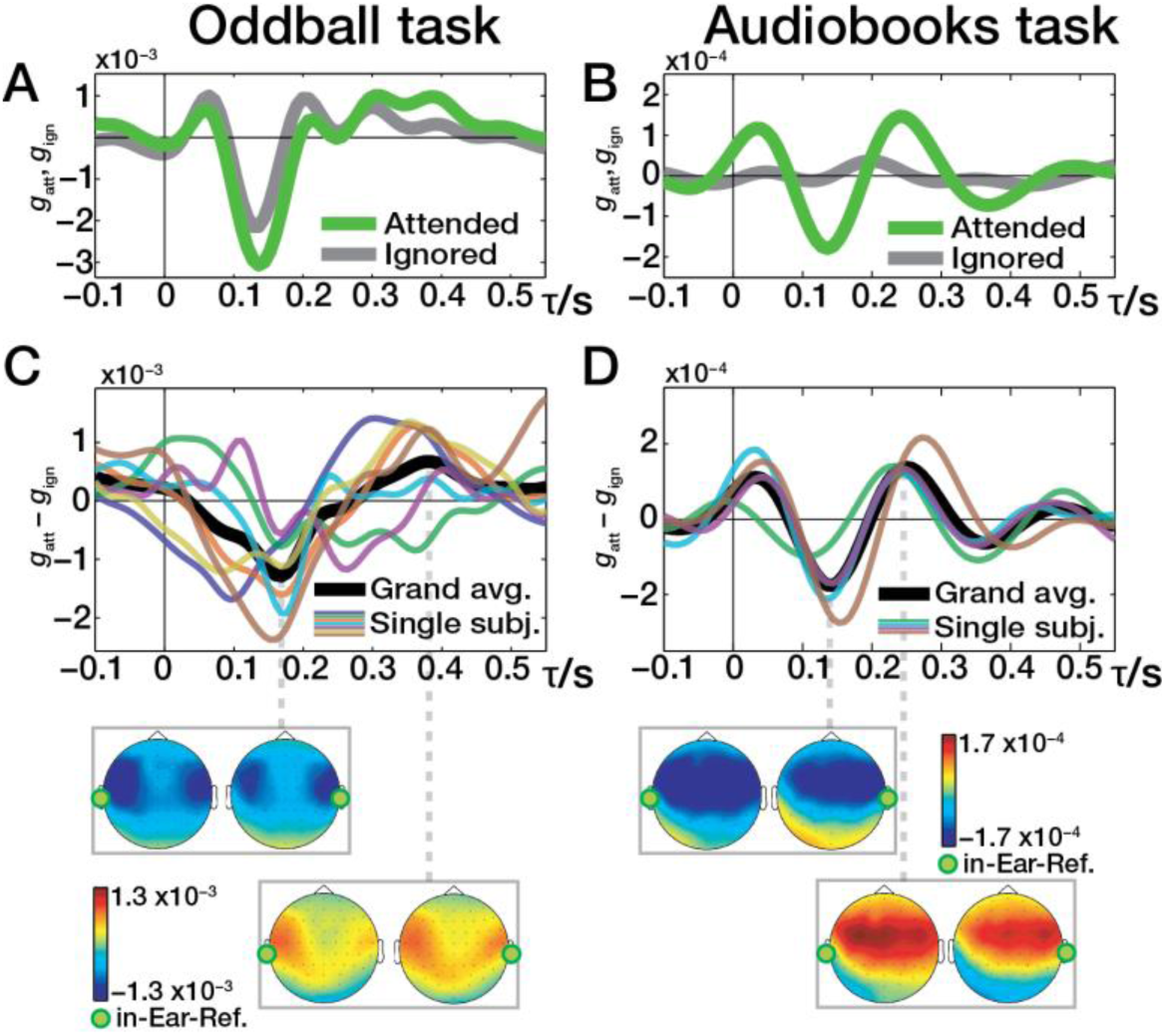
Response functions. Response functions shown here were obtained from potential difference between left in-Ear-EEG and FT7 electrode. **A)** Grand average response functions to both attended and ignored tones in the oddball task. **B)** Grand average response functions to both attended and ignored speaker in the audiobooks task. **C)** and **D)** show single subject data of difference between response functions in the oddball task and in the audiobooks task, respectively. Topographies show grand average weightings at time lags of maximal difference between the response functions (i.e., attended–ignored).

In the audiobooks task, a clear P50-N100-P200-equivalent complex could be found in the responses to the attended speaker (figure 3B). The responses to the ignored speaker show only weak magnitudes and suggest a suppression of the responses to the ignored speaker. Compared to the oddball task, this is leading to a greater difference between the responses to the attended and the ignored speaker (figure 3D). Again, the differences of the single subject’s response functions show a consistent pattern with a common negative deflection at a time lag of 130 ms and a later positive deflection at around 250 ms (figure 3D). The topographies of the components at 130 ms and 260 ms both have fronto-central patterns, spreading out towards temporal regions.

In both tasks, we have found response functions that show consistent patterns across subjects. In particular the deflections between responses to attended and ignored stimuli are prerequisites for a single channel classification approach (see above). Most interesting, these deflections could even be recorded at scalp-EEG electrodes located close to its in-Ear-EEG reference electrode.

### 3.2 Goodness of fit as a basis for identifying the attended stream

*Goodness of fit* was defined as correlation coefficient resulting from the Pearson-correlation of the measured EEG signal and the predicted EEG signal that consists of the responses to the to-be-attended and to-be-ignored stream (i.e., true prediction).

Generally, the average *goodness of fit* with values in a range of 0.02–0.15 (oddballs: mean = 0.12, range 0.08–0.15; audiobooks: mean = 0.04, range: 0.02–0.06) seems weak. In order to statistically evaluate if the correlations of the predicted and the measured EEG signals provide valuable information for classification, we investigated the distribution of the Fisher-z-transformed Pearson-correlation coefficients z_true_ and z_false_. Figure 4A & B show the distribution of the correlation coefficients in both tasks, where every single dot represents a single trial performed by a (colour-coded) single subject. The correlation of the true prediction and the measured EEG signal (z_true_) tends to be greater than its counterpart z_false_ in the majority of the trials (Figure 4A & B). The difference z_true_ — z_false_ was found to be significantly above zero for each subject (*one-sample t-test,* oddballs: six subjects *p < 0.001,* one subject *p < 0.01,* Figure 4C; audiobooks: two subjects *p < 0.001,* one subject *p < 0.01,* one subject *p < 0.05,* Figure 4D), suggesting it to be a valuable basis for deciding which of the streams is attended.

**Figure 4:**
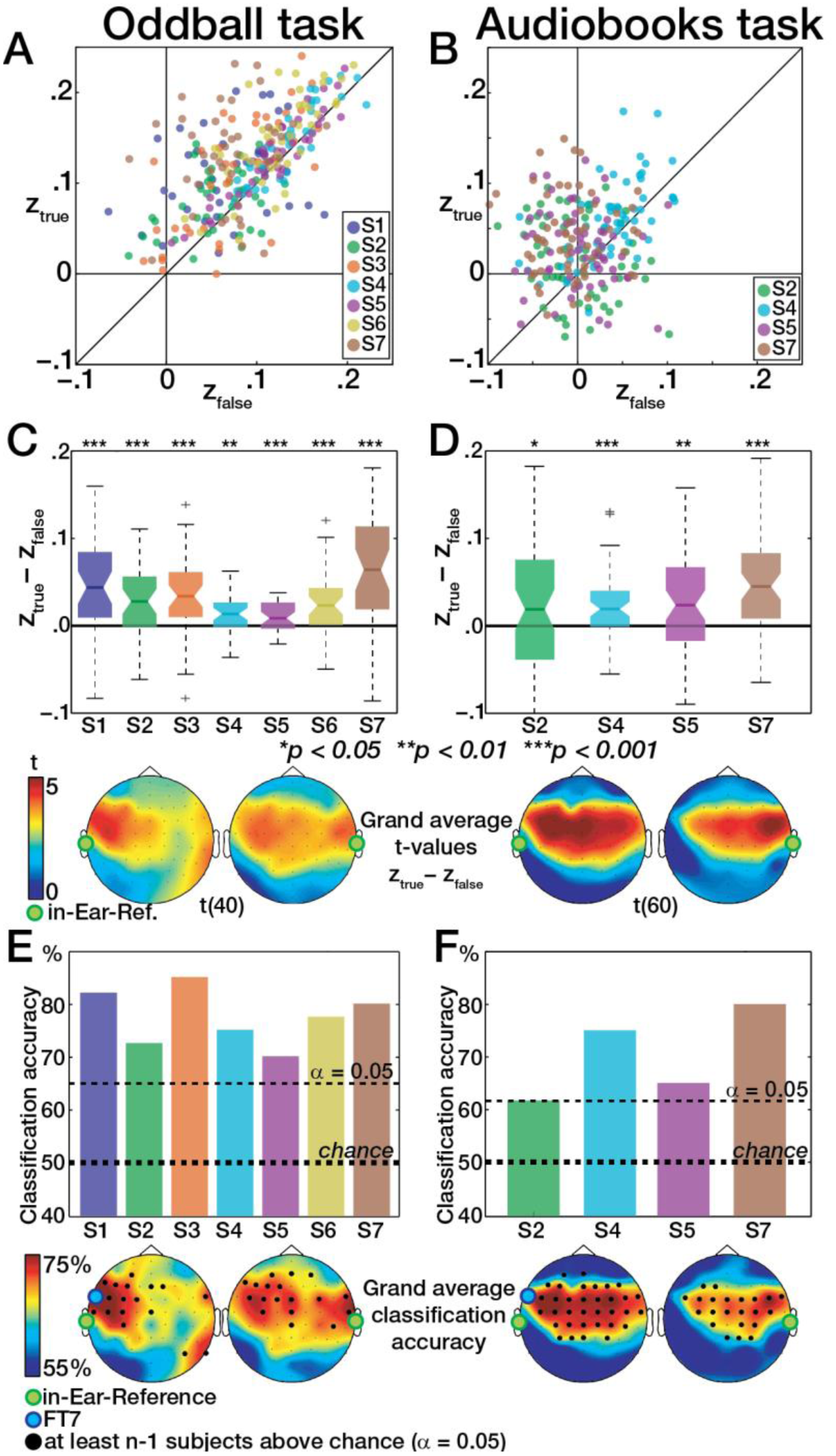
Goodness of fit and classification accuracy. Single subject data shown here were obtained from potential difference between left in-Ear-EEG and FT7 electrode. Topographies show grand average data. **A&B**) Each dot represents the relation of both Pearson-correlations z_true_ and z_false_ in single trials of the oddball task. **C&D)** Distributions of the difference z_true_ – z_false_ for single subjects, which were tested against zero (t-test). Topographies show grand average t-values. **E&F**) Classification accuracy based on the difference z_true_-z_false_. Horizontal lines indicate significance above chance based in a binominal distribution. Topographic maps show grand average classification accuracy. Highlighted channels are indicating channels where at least n-1 subjects yield classification accuracies significantly above chance.

In order to evaluate which electrode configuration provides best inference on identification of the attended speaker, we inspected the grand average topographies (Figure 4C & D) of the single subject t-values obtained from the distribution of the difference between z_true_ and z_false_ (see above). Strongest effects were found at in-Ear-EEG configurations incorporating fronto-central scalp-EEG channels. Interestingly, in both tasks highest t-values were observed for configurations consisting of scalp-EEG electrodes (i.e. FT7, FT8, T7, T8) close to the ear that the reference in-Ear-EEG electrode was placed in.

Generally, the analysis of *goodness of fit* gave insight how a set of two electrodes consisting of one electrode in the ear canal and another at the scalp close to the ear should be oriented in order to explain attention related variance in the EEG signal caused by auditory stimulation.

### 3.3 The attended stream can be identified from single-channel configurations

*Classification accuracy* was defined as the percentage of trials the predicted EEG signal labeled *true* yields a more positive Pearson-correlation coefficient with the measured EEG signal than the predicted EEG signal labeled *false.* For statistical analyses, Pearson-correlation coefficients were Fisher-z-transformed and called z_true_ and z_false_.

The classification accuracy at FT7 referenced to the left in-Ear-EEG electrode is shown in Figure 4E & F. Classification accuracy was found to be significantly above chance (p < 0.05) for all subjects and both the oddball task (mean: 77%, range 69–85%, Figure 4E) and the audiobooks task (mean: 70%, range 62-80%, Figure 4F) at this exemplary electrode configuration. With regard to the application in hearing aids, a purely in-Ear-EEG configuration consisting of two electrodes within the same ear canal is most desirable. We investigated those configurations as well and provided the results in the supplements (Figure S2). Note that these alternative configurations did not yield classification accuracy consistently above chance.

Grand average topographies of classification accuracy (Figure 4E & F) show patterns similar to the t-value topographies above (Figure 4C & D). Highlighted channels in Figure 4E & F indicate that classification accuracy was found to be above chance (p < 0.05) for at least all but one of the subjects. Interestingly, channels close to the ear the reference in-Ear-EEG electrode was placed in showed classification results above chance consistently across subjects.

Due to the low number of subjects, drawing a general conclusion on the most appropriate electrode configuration is not possible. However, for the present data we can state that we have found a configuration, showing classification results above chance for every subject consisting of only two electrodes, FT7 referenced to left in-Ear-EEG electrode. Single-subject topographical maps provided in the supplements (Figure S1 A) confirm that various short-distance electrode configurations yield classification accuracy above chance. Based on the single-channel data of subjects who participated in both tasks, we found a strong dependency of classification accuracy between tasks (Figure S1 B), which emphasizes the robustness of our findings despite our relatively low number of participants.

## 4 Discussion

It is a frequently stated long-term goal to fuse EEG recordings with hearing aid technology in order to attune the hearing aid to an attended sound source. Here, we investigated whether the attended sound stream out of two concurring streams can be identified from single channel EEG-recordings. Single channels were electrode configurations consisting of one reference in-Ear-EEG and one scalp-EEG electrode. We focused our analyses on a configuration consisting of a left in-Ear-EEG electrode and scalp-EEG electrode FT7.

Participants performed two tasks. In both tasks, concurrent sound streams (i.e. tones and speech) were presented. We hypothesized single channel in-Ear-EEG data to provide valuable information to identify the attended stream.

### 4.1 Response functions consistently reveal listeners’ focus of attention

In contrast to backward models, the estimation of forward models allows the comparison of the obtained response functions with conventional ERPs (Lalor *et al* 2009). An attention-related difference between response functions is a prerequisite for identification of the attended speaker (see Methods).

In both tasks, we have found an enhanced N100-equivalent component in the responses to attended stimuli compared with ignored stimuli for each subject (Figure 3A & B). This is in line with auditory evoked potential (AEP) studies, showing that the N100 component is enhanced if the stimulus is attended (e.g., Näätänen *et al* 1981).

Notably, attention-related differences in the response functions could be found even in short-distance configurations consisting of a reference in-Ear-EEG electrode and a scalp-EEG electrode close to the ear, as exemplarily shown for FT7 referenced to left in-Ear-EEG electrode. In regards to hearing aid applications, these findings encourage the attachment of only a few electrodes in the periphery of the ear (Mirkovic *et al* 2016).

The consistent morphology of the difference between responses to attended and to ignored stimuli (Figure 3C & D) further suggests the training of a model based on the data of all but one subject and test it on the latter (i.e., generic model). Even if not as accurate, O’Sullivan *et al* (2015) showed that a generic model still allows predicting the attentional focus. With respect to its application in hearing aids, a generic model could provide a default set of parameter values before a listener-specific model is adapted over time (Mirkovic *et al* 2015). In the current study, the training of a robust generic model was hindered by the low number of subjects and should be further investigated.

The dichotic oddball paradigm employed here also is appropriate when investigating neural responses to discrete and spatially separated stimuli. However, such a paradigm is removed from real-world listening scenarios, since two or more sound sources in natural environments are rarely separated in a dichotic fashion and are rarely as stationary regarding their rhythm and spectral content.

In contrast, the audiobooks paradigm with two diotically presented speakers represents a challenging listening situation and is more akin to realistic scenarios (also with respect to a listener’s goal, that is, following a sound source and comprehending what is being conveyed (Obleser 2014). Since no spatial information is contained in the audio signal, a ‘worst case’ scenario was presented. Sound source separation can only be achieved based on spectral-temporal cues of the two speakers. Since each participant attended to either the male or to the female voice in the same number of trials, the revealed differences of the response function can’t be explained by spatially separated stimuli nor from speaker specific features.

In most of the cited studies on detection of auditory attention from EEG data, the speech envelope was used as stimulus representation (O’Sullivan *et al* 2015, Mirkovic *et al* 2015, Biesmans *et al* 2016). In contrast we used onset envelopes, that is, the halfwave-rectified first derivative of the envelope. Using instead the envelope led to similar detection accuracies (Figure S4 A), but responses were shifted by approximately 50 ms such that the P50 equivalent component appeared before time lag of zero (Figure S4 B). This is due to every onset being followed by a peak in the envelope after approximately 50 ms (Figure S4 C). For the oddball task, the correct latencies of the components (i.e. P50, N100, P200) are known from previously calculated ERPs (Fiedler et al 2016). Since the latencies of the onset envelope responses in the audiobooks task fit the latencies of the ERP onset responses in the oddball task better than the envelope responses do, we conclude that onset envelopes lead to more precise estimations.

A comparison of the response functions reveals similar latencies of components between tasks, but the relative suppression of the response to the ignored stream is stronger in the audiobooks task. Two diotically presented speakers are more likely masking each other than dichotically presented tones of 100 ms length (and up to 614 ms pauses between tones). The suppression of the responses to the ignored speaker might indicate higher demand for suppression of the ignored stream and thus a higher task difficulty.

Of course, the low number of individually in-Ear-fitted subjects tested here (*n* = 7 & n = 4) allows only for limited conclusions. However, the markedly consistent morphologies of the response functions and the individually significant detection success suggest that differential responses to attended and ignored auditory stimuli, even continuous speech, can be recorded from short-distance electrode configurations. These configurations here consisted only of one electrode in the ear canal and another close to the same ear, as exemplarily shown in Figure 4E & F for a left in-Ear-EEG electrode referenced to scalp-EEG electrode FT7. Please note that the shortest distance we could achieve was determined by the electrode positions of the scalp EEG. The exemplary electrode FT7 is placed at a distance of approximately 8 cm to the entrance of the ear canal (tragus) at an angle of 40° relative to the tragus-Cz-line. With the development of adhesive electrodes to be attached around the ear it was shown that responses could be recorded at even closer positions (Bleichner *et al* 2016).

### 4.2 Goodness of fit provides basis for identification of the attended stream

Former studies about approaches to identification of the attended speaker mainly used backward decoding models (O’Sullivan *et al* 2015, Mirkovic *et al* 2015, 2016, Biesmans *et al* 2016). Backward models are trained on multi-channel EEG data and used to reconstruct a single speech envelope. In contrast, we used forward models to predict the EEG signal in response to the stimulus, which allowed us to quantify the *goodness of fit* at every single EEG channel (see Methods).

The *goodness of fit* was quantified by Pearson’s correlation-coefficient for the predicted versus the measured EEG signal. In the previous backward model studies cited above, correlation-coefficients obtained from Pearson-correlation of the reconstructed and the original speech envelope between 0.02 and 0.10 were reported. Here, we obtained correlation coefficients of similar magnitude, but they were here obtained solely on the basis of a potential difference recorded at a single EEG-channel consisting of left in-Ear-EEG and scalp-EEG electrode FT7. Crucially, the topographies of single-trial-derived t-values (Figure 3C & D) show that meaningful differences can be found satisfyingly at single electrodes close to the referenced in-Ear-EEG electrode.

We thus conclude that short-distance electrode configurations like the exemplary configuration consisting of the left in-Ear-EEG reference and FT7 electrode capture information about the listener’s attentional focus and thus provide a basis for the identification of the attended sound source. To achieve this, we based our analyses on certain assumptions. First, we assumed that strongest responses can be found at stimulus onsets and thus extracted respective representations (see Methods). Especially for speech, features known to evoke responses are manifold and rarely mutually exclusive, since all are, to some extent, nested or derived from the broad-band temporal envelope (Ding and Simon 2014). Second, we applied ridge regression in order to train a model under the assumption of linearity and with the goal to reduce the mean squared error of the prediction. The extraction of features from speech is wedded to the selection of an appropriate model and both affect the contrast between responses to attended and ignored speech.

Comparing several methods of extracting features of speech and going beyond the simple assumption of linearity as well as incorporating several loss-functions might further boost the contrast between the two predicted EEG signals and thus further refine the information about the attentional focus.

### 4.3 The attended stream can be identified from single-channel configurations

The major goal of this study was to identify the attended sound stream based on single-channel hearing aid-compatible EEG channel configurations. Considering that, classification accuracy is the most important measure to evaluate the performance of our approach of single channel classification.

As stated above, former studies have used backward models to bring in the advantage of having multiple EEG signals to reconstruct one single speech envelope. In order to reduce the number of channels, Mirkovic *et al* 2015 already applied an approach of recursive channel elimination. Starting from a grid of 96 channels, it was shown that a stepwise exclusion of worst performing channels doesn’t affect classification accuracy up until approximately 25 channels were left. The best performing electrodes were concentrated at temporal positions close to the ear. However, the average of all electrodes served as reference potential which hinders a conclusion for single channel configurations consisting of only two electrodes. In a recent study (Mirkovic *et al* 2016), it was shown that based on the data of a grid of ten electrodes around the ear the attended speaker could be identified with a backward model. Here, we go even further and show that a montage of only two electrodes, left in-Ear-EEG electrode and scalp-EEG-electrode FT7, is sufficient to identify the attended sound source in two experimental tasks. In Mirkovic *et al* (2016), we presume that placing a few electrodes at positions favorable for identifying the attended speaker is more crucial than obtaining more or less redundant EEG signals from multiple channels.

With respect to the long-term goal of controlling a hearing aid in real-time, our results provide valuable insight. First, in a hearing aid, computational resources are limited. We thus decided not to apply any method of artifact rejection or other methods of signal enhancement other than band-limiting the EEG-signal. Once a model is trained, the algorithm consists of only four convolutional operations and two correlations. Considering the comparably low sampling rate of 125 Hz and one-minute trials of 7500 samples, the computational effort is comparably low.

Nevertheless, a classification accuracy of around 70% after one minute might not yet comply with the requirements of a hearing-aid user. Furthermore, data were recorded in a shielded room which reduced environmental noise as well as subjects were asked to move as less as possible which lead to a minimum of muscle artifacts. Please note that an implementation of such an electrode configuration into a hearing-aid would raise further issues not addressed here, such as how to attach an electrode outside the ear canal and dealing with low conductance due to hairy positions and skin resistance. One possible solution might be permanently or daily placed electrodes around the ear (Debener *et al* 2015, Mirkovic *et al* 2016, Bleichner *et al* 2016). Thus, for real-life applications, there are still major challenges ahead. Our findings however do map out a significant step towards the application of single channel in-Ear-EEG in future hearing aids.

## 5 Conclusion

The identification of attended sound sources based on neural data has become increasingly important for both, neuro-scientists and hearing aid developers, since it contains the potential to control a hearing prosthesis in a brain-computer interface fashion. One unsolved problem is the embedding of EEG electrodes and utilization of EEG signals in the hearing-aid periphery.

In the current study, we have shown that in-Ear-EEG can feasibly capture information about the listeners’ attentional focus. Thus, with only two electrodes attached, an auditory brain-computer interface could constantly track a listener’s attentional focus. This information could be fed back to other hearing aid algorithms in real-time (e.g., controlling for directional microphones and noise suppression) at low computational cost.

## Acknowledgements

This work was supported by research grants from the VW Foundation (BIT-CHAT to JO and TL) and Oticon Foundation (NEURO-CHAT, to JO and TL). Thomas Lunner and Carina Graversen are with Eriksholm Research Centre A/S, part of Oticon.

## Supplements

**Figure S1:**
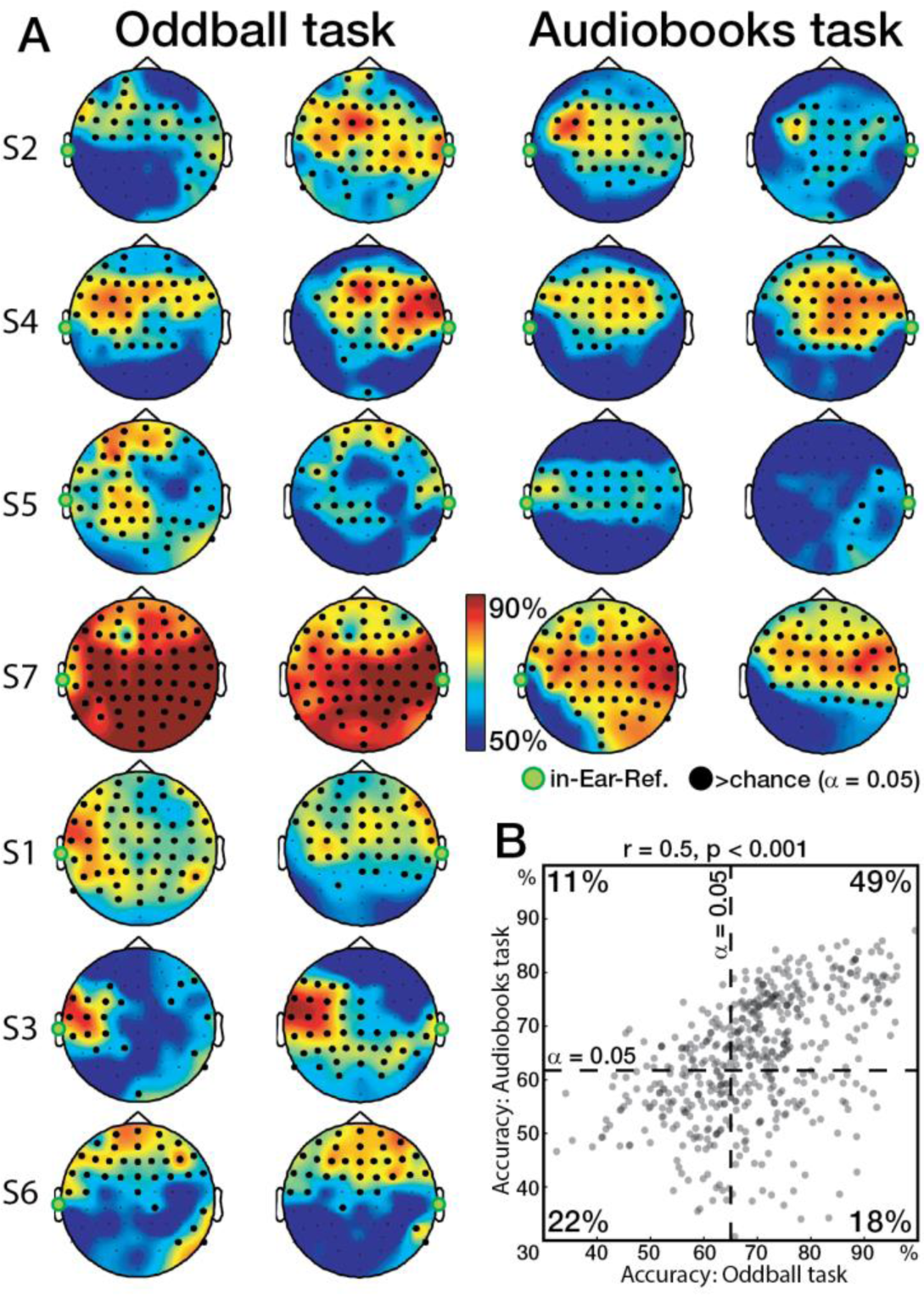
Single-subject classification accuracy. **A)** Single-subject topographical maps reveal similarity of patterns of classification accuracy between oddball and audiobooks task. **B)** Scatter plot of classification accuracy in both tasks at all single channels of all subjects who participated in both tasks (dots are slightly jittered to avoid overlap). Dashed lines indicate significance thresholds. The inspection of the number of dots falling into the quadrants revealed that channels either performing below chance in both paradigms (lower left quadrant) or above chance in both paradigms (upper right quadrant) make up 71%. This shows that we can estimate the classification accuracy of the audiobooks task from the classification accuracy of the oddball task. This finding is also supported by a significant (p < 0.001) Pearson-correlation coefficient of r = 0.5.

**Figure S2:**
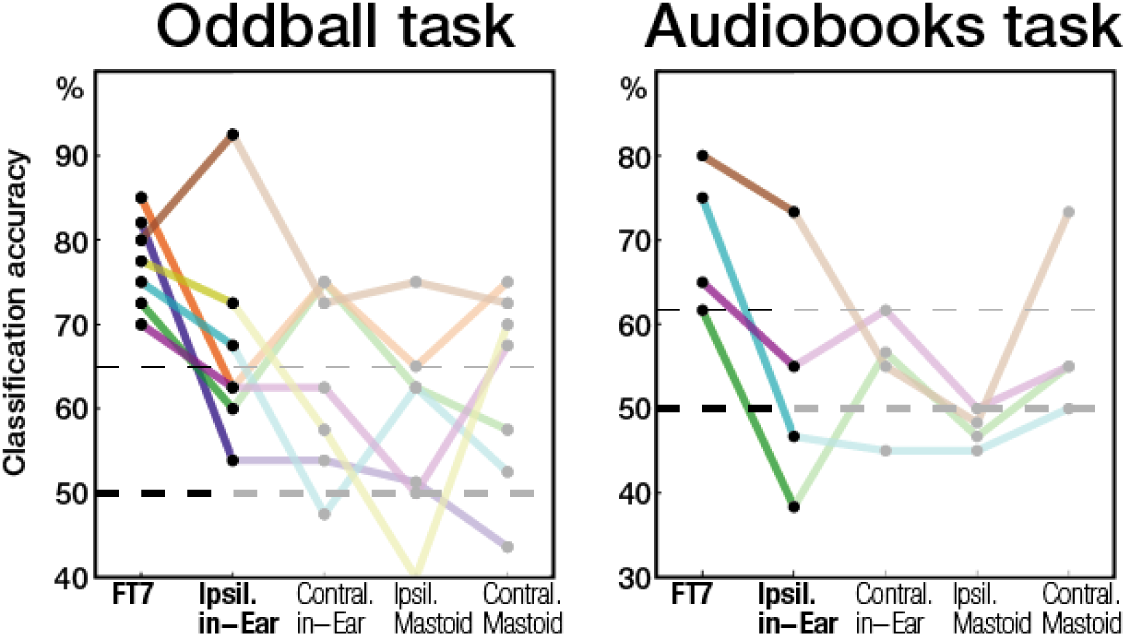
Comparison of single subjects’ classification accuracies at various electrode configurations. The left in-Ear-EEG electrode served as reference. With regard to the application in hearing aids, the most useful configuration would be ipsilateral (i.e., same side as reference) in-Ear. This configuration yielded relatively reduced classification accuracies in all but one subject. Note that the ipsilateral in-Ear configuration for some subjects still yielded a classification accuracy above chance. Furthermore, we investigated the contralateral in-Ear EEG electrode, the ipsi-and contralateral mastoid. At these configurations, no consistent classification accuracy was found.

**Figure S3:**
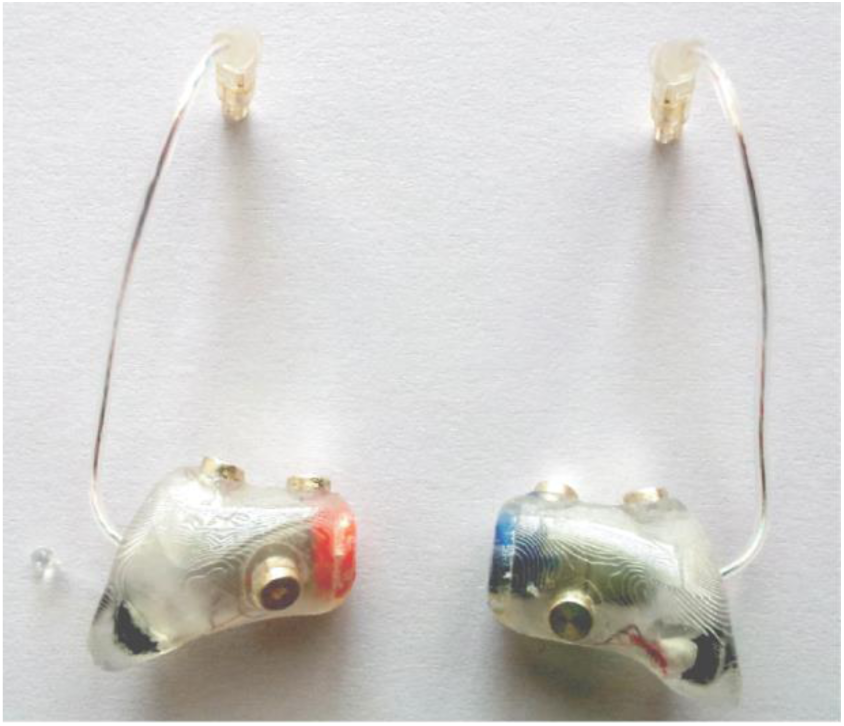
Exemplary in-Ear-EEG devices. Each device has three EEG electrodes attached (© 2016 IEEE. Reprinted, with permission, from Fiedler L, Obleser J, Lunner T and Graversen C 2016 Ear-EEG allows extraction of neural responses in challenging listening scenarios – a future technology for hearing aids? Eng. Med. Biol. Soc. (EMBC), 38th Annu. Int. Conf. IEEE, 38 5697–700). To create the in-Ear EEG devices, a trained audiologist took impressions of the ear canals, which were used to print 3D shells of each individual ear canal. Three holes were drilled in the shells to insert the electrodes, which were connected by short wires to a standard 3-pin Hi-PRO EASYFIT plug. Electrodes were made from a fine silverthread with a diameter of 3,00 mm cut into slices of 2,00 mm (Ravstedhus, Denmark).

**Figure S4:**
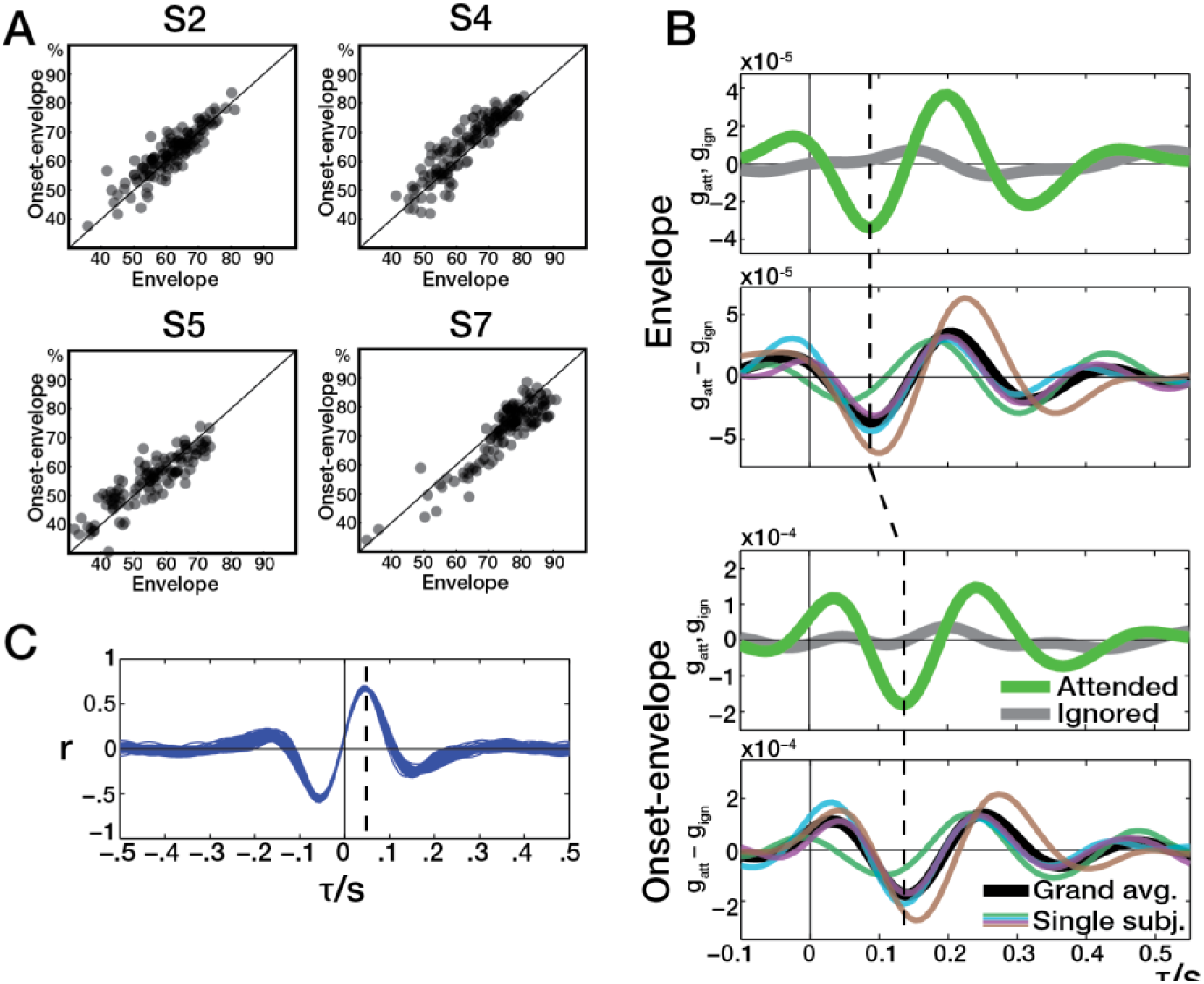
**A)** Relation between classification accuracies obtained with envelope and onset-envelope. **B**) Responses to envelope and responses to onset envelope. **C)** Cross-correlation of onset-envelope and envelope. Blue lines indicate individual trials. Positive time lags shift the onset-envelope towards the future, leading to maximum similarity with the envelope at a time lag of approximately 50 ms.

